# Multiomic profiling of medulloblastoma reveals subtype-specific targetable alterations at the proteome and N-glycan level

**DOI:** 10.1101/2023.01.09.523234

**Authors:** Hannah Voß, Shweta Godbole, Simon Schlumbohm, Yannis Schumann, Bojia Peng, Martin Mynarek, Stefan Rutkowski, Matthias Dottermusch, Mario M. Dorostkar, Andrey Koshunov, Thomas Mair, Stefan M. Pfister, Philipp Neumann, Christian Hartmann, Joachim Weis, Friederike Liesche-Starnecker, Yudong Guan, Hartmut Schlüter, Ulrich Schüller, Christoph Krisp, Julia E. Neumann

## Abstract

Medulloblastomas (MBs) are malignant pediatric brain tumors that are molecularly and clinically very heterogenous. To unravel phenotypically relevant MB subtypes, we compiled a harmonized proteome dataset of 167 MBs and integrated findings with DNA methylation and N-glycome data. Six proteome MB subtypes emerged, that could be assigned to two main molecular programs: transcription/translation (pSHHt, pWNT and pGroup3-Myc), and synapses/immunological processes (pSHHs, pGroup3 and pGroup4). Multiomic analysis revealed different conservation levels of proteome features across MB subtypes at the DNA-methylation level. Aggressive pGroup3-Myc MBs and favorable pWNT MBs were most similar in cluster hierarchies concerning overall proteome patterns but showed different protein abundances of the vincristine resistance associated multiprotein complex TriC/CCT and of N-glycan turnover associated factors. The N-glycome reflected proteome subtypes and complex-bisecting N-glycans characterized pGroup3-Myc tumors. Our results shed light on new targetable alterations in MB and set a foundation for potential immunotherapies targeting glycan structures.

**Significance:** Whereas the application of omics technologies has significantly improved MB tumor classification and treatment stratification, it is still of debate, which features predict best clinical outcome. Moreover, treatment options - especially for high-risk groups - are still unsatisfactory. In contrast to nucleic acids, the proteome and their N-glycans may reflect the phenotype of a tumor in a more direct way and thus hold the potential to discover clinically relevant phenotypes and potentially targetable pathways. We show that these analyses are feasible on formalin fixed and paraffine embedded tissue. Compiling a comprehensive MB dataset, we detected new biomarkers and characteristics for high- and low-risk MB subtypes that were not reflected by other omic data modalities before. Specifically, we identified subtype specific abundance differences in proteins of the vincristine resistance associated multiprotein complex TriC/CCT and in proteins involved in N-glycan turnover. Changes in the N-glycans are considered as potential hallmarks of cancer and we show that N-glycan profiles can distinguish MB subtypes. These tumor-specific N-glycan structures hold a strong potential as new biomarkers, as well as immunotherapy targets.

**Highlights:**

- Integration of in-house proteome data on formalin fixated paraffine embedded medulloblastoma (MB) and publicly available datasets enables large scale proteome analysis of MB
- Six proteome MB subtypes can be assigned to two main molecular programs: replication/ translation versus synapse/immune system
- Identification and validation of IHC compatible protein-biomarkers for high and low risk MB subtypes, such as TNC and PALMD.
- Subtype specific correlation of the DNA methylome and the proteome reveals different conserved molecular characteristics across MB subtypes.
- pGroup3-Myc subtype MBs are associated with high-risk features including high abundances of vincristine resistance associated TriC/CCT member proteins
- Proteome MB subtypes show differential N-glycosylation patterns, revealing complex-bisecting glycans as potentially immunotargetable hallmarks of the high risk pGroup3-Myc subtype.

## Introduction

Medulloblastomas (MBs) are aggressive pediatric brain tumors, occurring in the posterior fossa. Histomorphologically, molecularly and clinically, MBs are a heterogenous disease, which is recognized by the current classification system (WHO)^1^. Four main consensus subtypes have been described, WNT pathway activated MB (WNT MB), Sonic hedgehog (SHH) pathway activated MB (SHH MB), and Group 3 (G3) and Group 4 (G4) MB^2^. Advances in molecular analyses, mainly using gene expression profiling, next generation sequencing and DNA methylation analysis, led to the discovery of substantial intragroup MB heterogeneity and many different subtypes of MBs with associations to distinct clinical features were described^3–6^. Certain markers for poor survival have been identified among MB subtypes, e.g. anaplastic histology, *MYC* amplification status, methylation subtype II/III, or *TP53* mutations in WNT and SHHMB^7–12^. Conversely, extensive nodularity (MBEN histology) and WNT activation (e.g. nuclear accumulation of β-CATENIN or CTNNB1 mutations) were associated with a favorable prognosis in MB patients ^12–14^. Additionally, methylation subtype VII or a distinct whole chromosomal alteration signature in non-WNT/ non-SHH MB was described as a predictor for a favorable outcome^12,15,16^. Whereas the clinical association to certain methylation subtypes and chromosomal aberrations has been clearly described, the underlying molecular mechanisms remain to be resolved and targeted treatment options are still lacking. In contrast to nucleic acids, the proteome and post-translational modifications may reflect a tumor’s phenotype in a more direct way and hold the potential to more precisely dissect clinically relevant phenotypes and targetable functional alterations. Furthermore, diverse genomic or epigenomic alterations might merge to similar proteomic patterns that could reveal common therapeutic targets among MBs. Studies on small MB cohorts, using fresh frozen (FF) tumor material, have shown that MBs also display heterogeneity at the proteome level ^17–19^. Formalin fixed and paraffine embedded (FFPE) material, enables the generation of larger datasets with higher statistical power but provides challenges to mass spectrometric analysis due to the incomplete reversion of methylene bridges and the induction of irreversible chemical modifications^20^.

In addition to the general protein abundance distribution, post-translational modifications (PTM) of proteins are important to understand cell physiology and disease related signaling networks^21^. Differential protein phosphorylations and acetylations have been described in the brain tumor context^17–19^. The most complex and common PTM, N-glycosylation, has not been targeted in MB yet. Changes in the N-glycome are considered potential hallmarks of cancer and N-glycan structures hold a strong potential as new biomarkers, as well as immunotherapy targets^22–26^.

Using a missing-value tolerant pipeline for integrating proteome data^27^, we incorporated publicly available, smaller MB proteome datasets^17–19^ with data of 62 FFPE MB cases and established a joint MB proteome dataset (n = 176). We comprehensively correlated proteome data with DNA methylation data – a current gold standard for molecular brain tumor classification^28^. Targeting N-glycosylation patterns across MB subtypes, we additionally performed a global N-glycan analysis and correlated N-glycan information with disclosed proteome subtypes. Taken together we present a large integrated study of the MB proteome, DNA-methylome and N-glycome, revealing new insights into MB phenotypes, potential new biomarkers and therapeutic targets.

## Results

### Integration of in-house proteome data on FFPE MB and publicly available datasets enables large scale proteome analysis of MB

We analyzed 62 FFPE MB tumors (53 primary cases and 9 recurrences), using high-resolution liquid chromatography-tandem mass spectrometry (LC-MS/MS). Of these, 53 cases were additionally profiled using DNA methylation analysis. Missing value tolerant, nonlinear iterative partial least squares (NIPALS) principal component analysis (PCA) revealed a distinguishability of the four main molecular subgroups of MB (SHH, WNT, G3, G4)^2^, that was previously described for fresh frozen (FF) tissue^17–19^ (Figure 1A, Supplementary Figures 1A, 2A). The age of used paraffine material did not impact on sample clustering (Figure 1B, Supplementary Figure 1A). Similar protein numbers and abundance levels of housekeeping proteins^29^ were obtained from FFPE material, generated over a span of 50 years (Supplementary Figure 1B, 1C). In addition, proteins detected in WNT and SHH MB, showed similar tendencies in FFPE- and FF-MB datasets^18,19^ (Supplementary Figure 2B, C). We concluded that proteome analyses of FFPE MB tissue shows protein abundance distributions that are comparable to FF tissue derived analysis. To further increase the cohort size, we integrated protein abundances with reanalyzed LC-MS/MS data of FF-MB from public repositories^18^;^19^;^17^, Figure 1D-F). Missing value tolerant data harmonization ^27^ adjusted sample specific mean and CV values across studies (Supplementary Figure 3A) and enabled a clear separation of MB groups. (Figure 1E-F). Established protein biomarkers for molecular MB subtypes that were present in MS data, showed expected subgroup specific abundance patterns (GAB1, CTNNB1, FLNA^30^, Figure 1G). Abundance levels of the housekeeping proteins were constant across integrated datasets (Supplementary Figure 3B). In total 16,279 proteins were quantified across 167 samples from the integrated dataset (19xWNT; 57xSHH; 53xG4; 36xG3; 2xno initial main subgroup stated), including 156 primary tumors and 11 recurrences.

**Figure 1:**
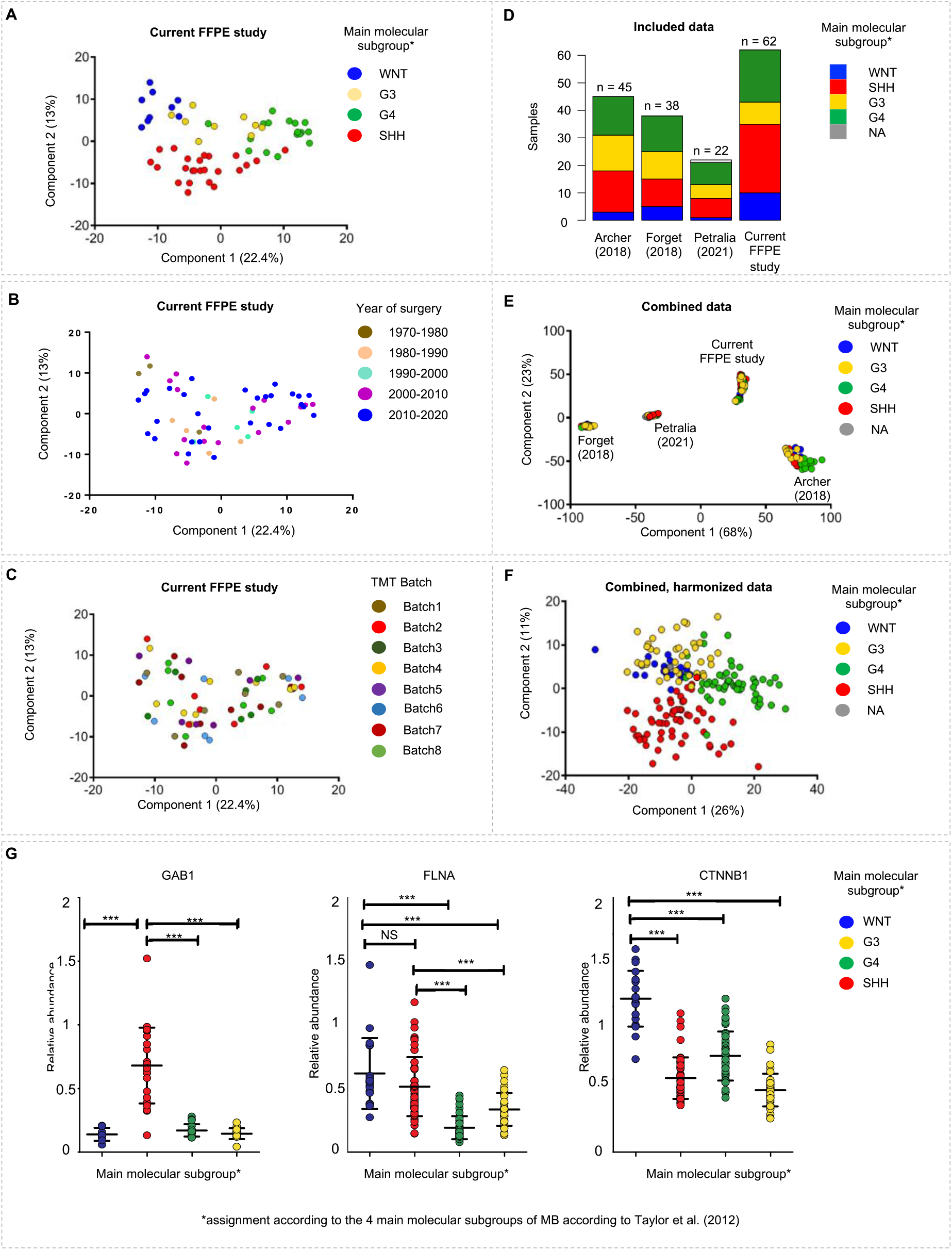
Harmonization and integration of proteome Medulloblastoma (MB) datasets. NIPALS principal component analyses (PCA) of measured FFPE samples (n=62) with assignment to **(A)** the four main molecular MB subgroups^2^, **(B)** age of measured samples, **(C)** measured MT batch. **(D)** Overview of datasets. NIPALS PCA of data before **(E)** and after **(F)** data harmonization using ComBat in the HarmonizR framework (n=167). **(G)** Protein abundance of the SHH MB marker GAB1, the WNT and SHH MB marker FILAMIN A and the WNT MB marker CTNNB1 after harmonization. *: p<0.05, **p<0.01, ***p<0.001

### Six proteomic MB subtypes can be assigned to two main molecular profiles

We next asked how many subtypes of MB are reflected at the proteome level. Consensus clustering, using hierarchical-and k-medoids (PAM)-clustering with different measures (Pearson, Spearman, Euclidean distance) identified 6 stable clusters, confirming previously described results of a smaller MB cohort (Figure 2 A-D)^18^. Proteome clusters partly overlapped with previously described subgroups, based on DNA methylation (Figure 2D, Supplementary Table 1b)(https://www.molecularneuropathology.org/mnp/)^28^. The assignment reliability of a sample to a respective proteome subtype was indicated by how many times a sample was associated with a certain cluster (cluster certainty, Figure 2D).

**Figure 2:**
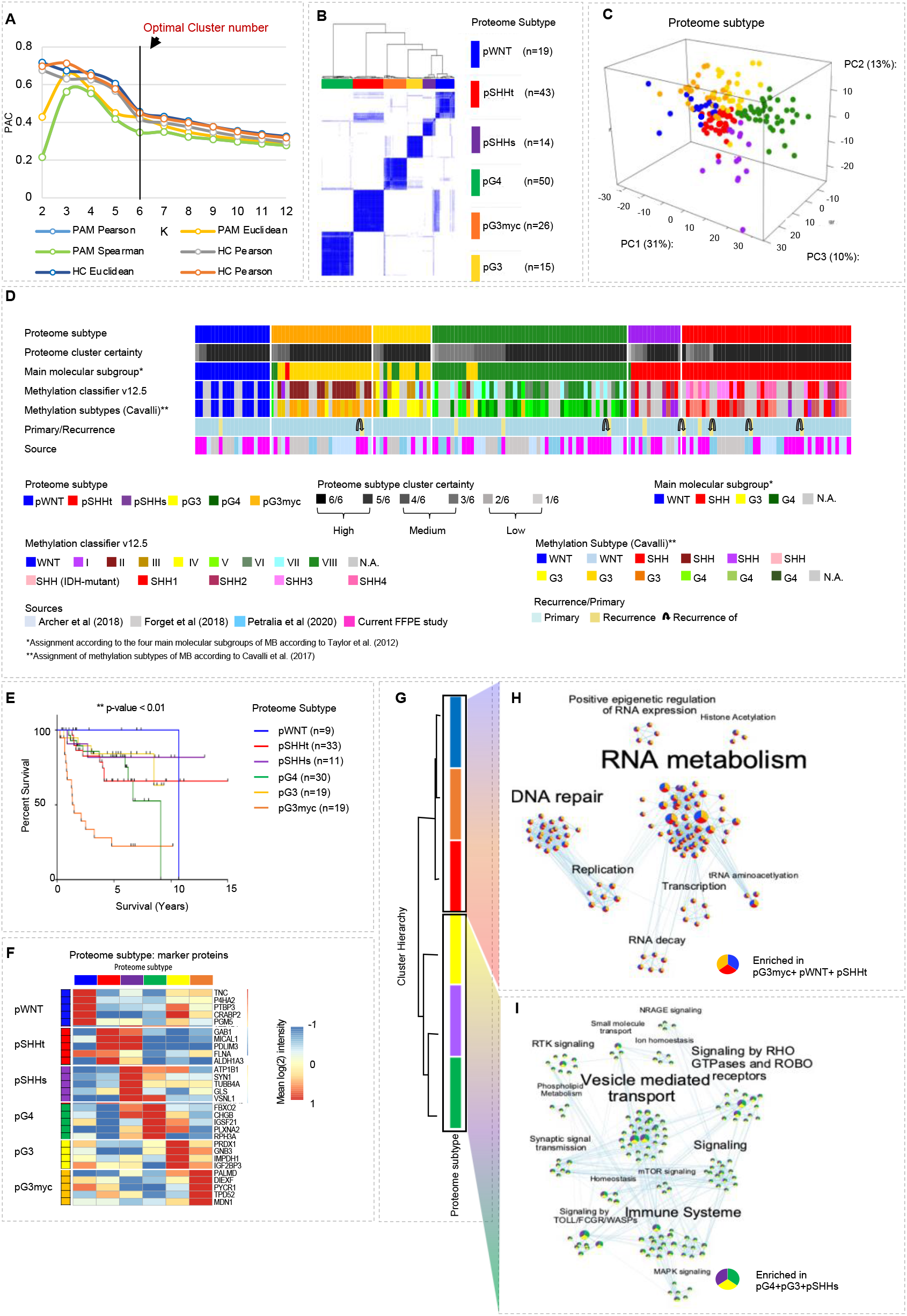
MB segregate into six proteome subtypes. **(A)** Proportion of ambiguous clustering (PAC) scores for k=2-12 in consensus clustering, using different cluster algorithms (n_MB_ = 167). **(B)** Optimal clustering of proteome data. Consensus scores shown in color scale from white (samples never cluster together) to blue (samples always cluster together). Six proteome subtypes, pWNT, pSHH-t, pSHH-s, pGroup3-Myc, pGroup3 and pGroup4, were defined. **(C)** Visualization of the first three principal components from NIPALS PCA. **(D)** Clinical sample information. (**E)** Log-rank (Mantel-Cox) test comparing the survival curves of proteome subtypes (p value < 0.001, overallχ2-square test). **(F)** Group specific mean log 2 protein intensity of protein subtype marker candidate proteins. **(G)** Proteome cluster similarity hierarchy based on stepwise increasing k-means execution from k=2-6. **(H)** Gene set description-based, gene set overlap dependent MCL clustering of enriched gene sets, comparing pWNT, pG3myc, pSHHt (n=88) to pG3, pG4 and pSHHs (n=79), Gene set enrichment analysis (GSEA) was based on REACTOME pathways for all analysis.

At the proteome level, non-WNT/ non-SHH MBs divided into three groups (pG4, pG3myc and pG3, p=proteome group), while SHH MBs separated into two proteome groups (pSHHs, pSHHt, s=synaptic profile, t= transcriptional profile). WNT MB formed a homogenous cluster (pWNT, Figure 2D). In general, a high cluster stability was given for all proteome subtypes (median 6/6), with the lowest mean cluster certainty given for pG3 samples, as they showed high similarity to pG4 and pG3myc respectively (median pG3 5/6, Figure 2D).

Proteome MB subtypes were associated with DNA methylation subtypes^5,6,31^ (https://www.molecularneuropathology.org/mnp/), Figure 2D, Supplementary Figure 4, Supplementary Figure 5, details see below). Except for one case, corresponding recurrent and primary tumors were assigned to the same proteome subtype (Figure 2D). The case that switched subtype in the recurrence situation (from pSHHs to pSHHt) had a low cluster certainty in the primary sample (3/6, Figure 2D).

pG3myc patients showed a significantly reduced overall survival as expected for subtype II/ *MYC* amplified samples (Figure 2E). pWNT patients, as expected, showed the best overall survival rate (Figure 2E). To determine potential biomarkers for proteome subtypes of MB, Students t-testing was performed. In total, 529 out of 3,996 proteins, found in at least 30% of all samples for each proteome subtype, showed a significant differential abundance in at least one subtype. For each group, the top 5 proteins with the lowest p-value and highest mean difference were selected as potential biomarker candidates (Figure 2F, Supplementary Table 2a). For high-risk non-WNT/non-SHH MBs (pG3myc) PALMD, DIEXF, MCN1 and PYCR1 were identified as potential new biomarkers, alongside with the previously described prognostic marker TPD52^8^. Of note, hedgehog-signaling induced proteins (MICAL1, GAB1, PDLIM3)^32^ showed a higher abundance in both, pSHHt and pSHHs. Subtype assignments and biomarker profiles were confirmed in an additional proteome MB dataset^33^, including 5 specimens, that were previously analyzed in our integrated cohort^19^ (Supplementary Table 4j, k).

To determine the degree of similarity between proteome MB subtypes of the main dataset, consensus clustering was performed with k = 2 to k = 6, analyzing at which cluster number the above defined subtypes separate. The highest similarity was observed between pWNT and pG3myc as well as pG4 and pSHHs (Figure 2G). The clearly distinct clusters at k = 2 indicate that MB subtypes can be divided into 2 different molecular profiles at a superordinate hierarchy level of proteome subtypes: profile 1, including pWNT, pG3myc and pSHHt and profile 2, including pG3, pG4 and pSHHs (Figure 2H, I). Gene set enrichment analysis (GSEA) revealed 133 enriched REACTOME pathways in profile 2 (q-value <0.05, Supplementary table 2c). GSEA -based Markov Clustering (MCL), identified mainly synaptic and immunological processes and phospholipid signaling (Figure 2I). For profile 1, a replicative/transcriptional signature was observed (92 enriched gene sets, Figure 2H, Supplementary Table 2b). Taken together, 6 MB proteome subtypes of MB were detected in an integrated cohort, that could be assigned to two main molecular programs.

### Group specific correlation of the DNA methylome and the proteome reveals different conservation levels of molecular characteristics across proteomic MB subtypes

To get a better insight into the correlation of methylome and proteome data, multiblock data integration using sparse variant partial least squares discriminant analysis (sPLS-DA) was performed between DNA methylation data (115 samples, 10,000 differentially methylated CpG sites between the MNP v12.5 defined subtypes, Supplementary Figure 5 C,D, Supplementary Table 1d) and proteome data (115 samples, 3,990 quantified proteins present in 30% of samples)^34^. Among the most discriminative features for the defined groups, only a fraction, discriminating mainly the WNT subtype showed a correlation of proteome and DNA-methylation data (Figure 3A, arrows, Supplementary Figure 6, correlation cut-off > 0.7, Supplementary Table 3h). To refrain from any data bias, (such as feature extraction as just described), we next performed a MB subtype specific correlation between complete DNA methylome data (115 samples and 381,717 CpG sites) and proteome data (115 samples, 3,990 proteins, Figure 3B, C). While the number of proteins correlating with any random CpG site was high and relatively similar for all MB subtypes (pWNT: 3,980 proteins, pG3: 3,990 proteins, pG3myc: 3,990 proteins, pG4: 3,340 proteins, pSHHs: 3,977 proteins, pSHHt 3,926 proteins), a significantly higher number of proteins of the pWNT subtype (38.14%, 1,552 proteins) and pG3 subtype (45.41%, 1,812 proteins) correlated with at least one CpG site of their own gene when compared to the other subtypes (pG4: 1.92%, 77 proteins, pG3myc: 6.49%, 259 proteins, pSHHt: 4.68%, 194 proteins and pSHHs: 1.52%, 65 proteins, Figure 3B, Supplementary Table 3b-g). Only 12.2 % - 18% of the correlating CpG sites among subtypes were located at the transcriptional start site (TSS200, TSS1500, Exon1) of their respective gene (Figure 3B). To test the correlation for subtype specific biomarkers between protein and CpG site methylation, non-group specific correlation (Pearson correlation cut-off > 0.7) was performed (Figure 3C, Supplementary Table 3a). Out of 31 potential protein biomarkers (Figure 2F), 10 correlated with CpG sites of their own gene (Figure 3C, 3D). In summary, DNA-methylation changes assessed with the Illumina arrays were only partly reflected at the protein level, with highest conserved features among pWNT and pG3 subtypes between the proteome and methylome level.

**Figure 3:**
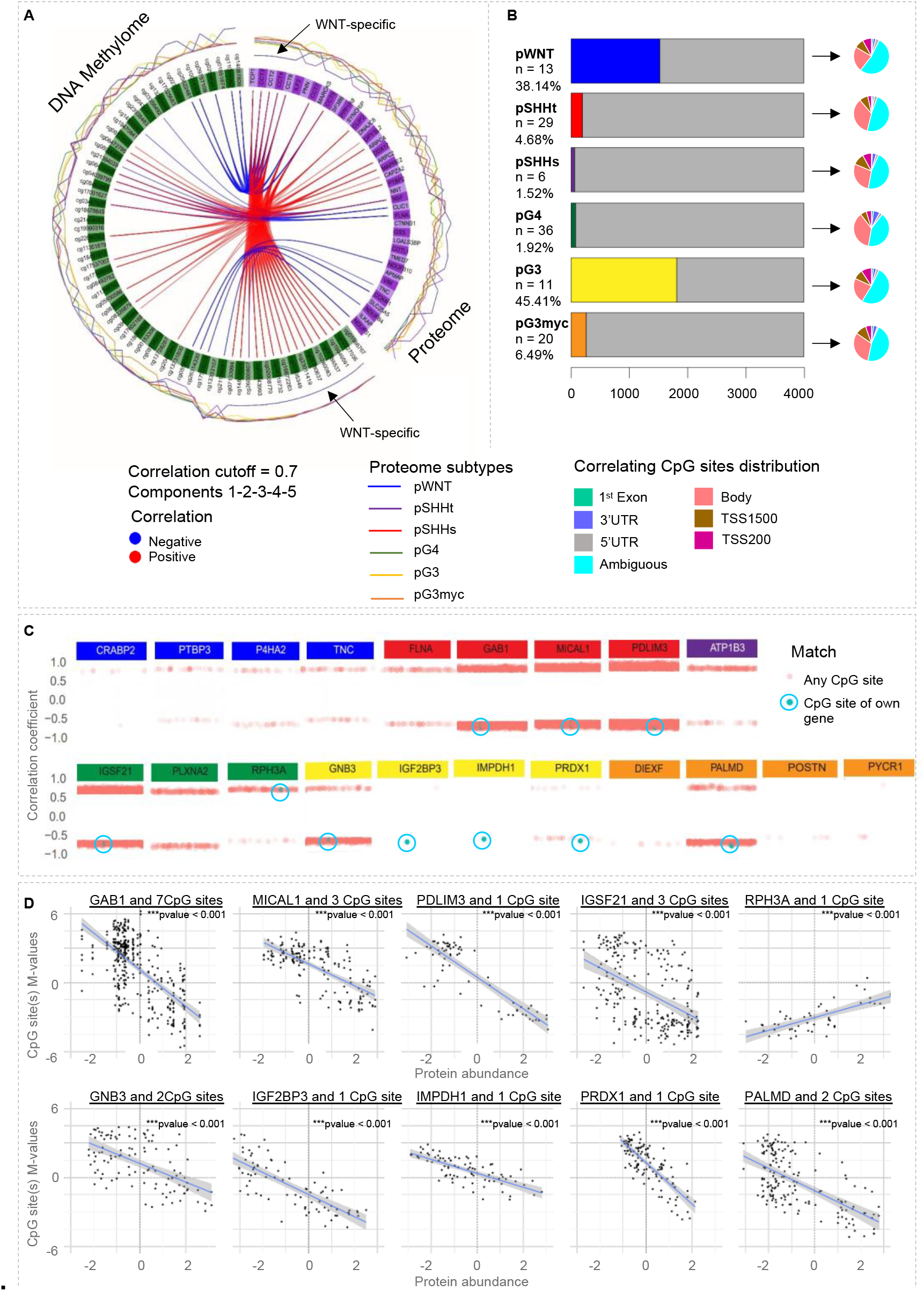
Correlation between DNA methylome and proteome features. **(A)** Circus plot from mixOmics analyses based on selected features of the first five components from proteome and methylome data. The plot illustrates features with correlation r > 0.7 represented on side quadrants. Proteome group specific feature levels are shown in the outer circle. **(B)** Proteome subtype specific Pearson correlation calculated between matched proteins and CpG methylation sites. The number of proteins correlating with CpG site methylation of their own gene (r > 0.7) is shown in colour. The pie chart shows the distribution of correlating CpG sites concerning the position in a gene. **(C)** Subtype independent Pearson correlation between 3,990 proteins and 381,717 methylation probes focusing on subtype specific biomarkers. Correlations >0.7 are shown, CpG sites correlating with the corresponding gene are highlighted in blue. Some biomarkers correlated with more than one CpG site of their own gene (GAB1: 7, GNB3: 2, IGSF21: 3, MICAL1: 3, and PALMD: 2). **(D)** Scatterplot of the 10 biomarker proteins correlating with the CpG site(s) of their own gene (correlation > 0.7, p < 0.001). The regression line was aligned for all correlating CpG site(s).

### SHH MB comprise two proteome subtypes showing either a synaptic or DNA transcription/translation signature

SHH MB splitted into two different proteome subtypes (pSHHt and pSHHs, Figure 4A). pSHHt, and pSHHs did not significantly correlate with histology or patient age. However, all pSHHs cases with high cluster certainty (6/6) occurred in patients below 3 years of age. The DNA methylation subtypes SHH3 (8/29) and SHH4 (9/29) were exclusively found in pSHHt MBs (Figure 4A). Methylation subtypes SHH1 and SHH2 were seen in pSHHs and pSHHt without statistical difference (SHH1: p=0.43, SHH2: p=0.10, X^2^ - test). SHH pathway alterations are regarded as driver events in SHH MBs^35^. *PTCH1* mutations were found exclusively but not mandatory in pSHHt tumors. *SUFU* and *SMO* mutations as well as amplifications of *MYCN* or *GLI2* did not distribute differentially (Figure 4A). *TP53* mutations are used for the stratification of high-risk SHH MB patients^36^. 9/10 *TP53* mutations occurred in the pSHHt subtype, but no significant differential distribution among pSHHt and pSHHs MBs could be verified (p= 0.43, X^2^ - test). The proteome abundance for each gene was mapped to chromosomal arms, which will be referred to as “proteome copy number variation (CNV)” henceforth. Both pSHHt and pSHHs groups showed a low overall correlation between calculated CNVs using DNA methylation data and proteome data (r_pSHHs_ =0.01, r_pSHHt_ =0.20, Figure 4D, G, Supplementary Tables 4g-h).

**Figure 4:**
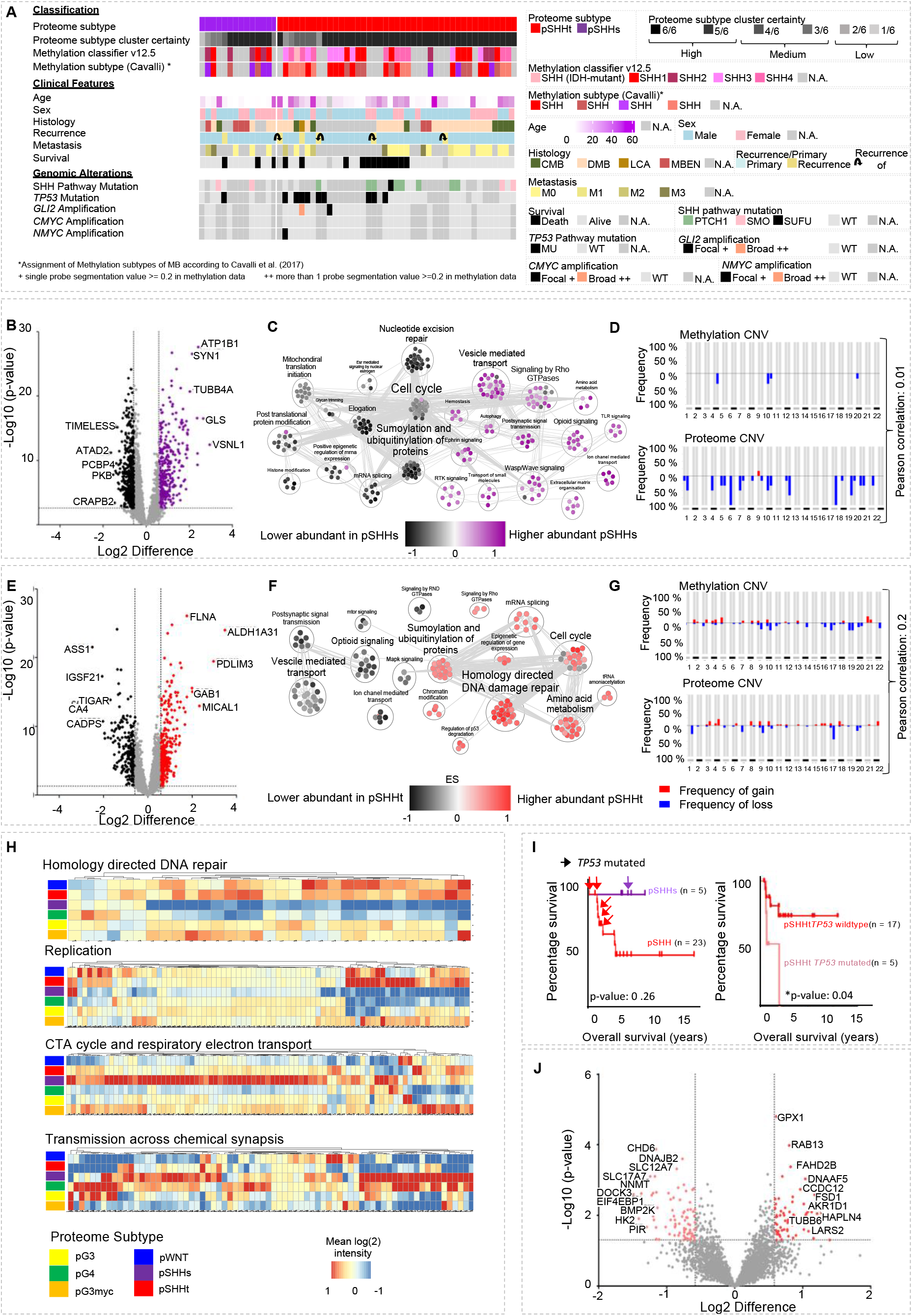
SHH MB comprise two proteome MB subtypes. ***(A)** Histological, molecular, and clinical characteristics of the MB subtypes pSHHt (n=43) and pSHHs (n=14). **(B)** Volcano plot showing differentially abundant proteins comparing pSHHs tumors to all other proteome subtypes. (p-value<0.05; Foldchange difference > 1.5). **(C)** GSEA based* Markov Cluster *Algorithm (MCL) clustering enriched gene sets in pSHHs MBs. **(D)** Copy number variations (CNV) plots of pSHHs MB (n=6) calculated from either DNA methylation or proteome data with correlation between both omic types (Pearson, r=0.01). **(E)** Volcano plot showing differentially abundant proteins when comparing pSHHt tumors to all other proteome subtypes (p-value<0.05; Foldchange difference > 1.5). **(F)** Gene set overlap dependent MCL clustering of enriched gene sets in pSHHt. **(G)** CNV plots for pSHHt MBs (n= 29) calculated from either DNA methylation or proteome data with correlation between both omic types (Pearson, r=0.2) **(H)** Heatmaps showing mean MB subtype protein abundance hallmark genesets homology directed repair (NES_pSHHt_= 2.2, p= <0.0001), replication (NES_pSHHt_= 2.2, p= 0.01), CTA cycle and respiratory electron transport (NES_pSHHs_= 3.9, p = <0.0001) and transmission across chemical synapses (NES_pSHHs_= 3.2, p = <0.0001). **(I)** Overall survival of pSHHt MB (n= 23) and pSHHs MB (n=5) and overall survival of pSHHt MB depended on TP53 mutation status. TP53 mutated cases displayed a significantly worse survival (Mantel cox test p-value = 0.04). **(J)** Volcano plot, showing differentially abundant proteins when comparing TP53 mutated cases to wildtype cases in pSHHt tumors (p-value<0.05; foldchange difference > 1.5)*.

pSHHs showed 167 differentially abundant proteins (Figure 4B, Supplementary table 4a). GSEA revealed 80 enriched REACTOME pathways associated mainly with synaptic, mitochondrial and immunological processes (q-value < 0.05, Figure 4C, Supplementary table 4c-d). 131 differentially abundant proteins detected in pSHHt showed an enrichment of post translational protein modification, transcription/translation, DNA repair and cell cycle associated gene sets (Figure 4E, 4F, H, Supplementary Table 4b, e-f). Of note, gene sets involved in “Signaling by RHO GTPases” and “Amino acid metabolism”, were enriched in both proteome SHH MB subtypes. The synaptic signature of pSHHs resembled the protein abundance distribution of “Transmission across chemical synapses” related proteins in pG4 (Figure 4H, see also Figure 5B). In contrast, the high abundance of mitochondrial proteins was exclusively found in pSHHs (Figure 4H). As expected, *TP53* mutations, assigned to the pSHHt group significantly correlated with bad prognosis in SHH patients (Figure 4I), while no significant survival difference was identified between pSHHs and pSHHt (Figure 4I). Based on the highly abundant proteins used for detection of statistically valid biomarkers for each of six proteome subtypes, *TP53* mutations did not lead to the formation of a distinct proteome cluster. However, when comparing *TP53* mutated with *TP53* wildtype MB within the pSHHt subtype, 134 proteins showed a statistically significant differential abundance between pSHHt-*TP53* wildtype and pSHHs-*TP53* mutated (Figure 4J, Supplementary Table 4i).

**Figure 5:**
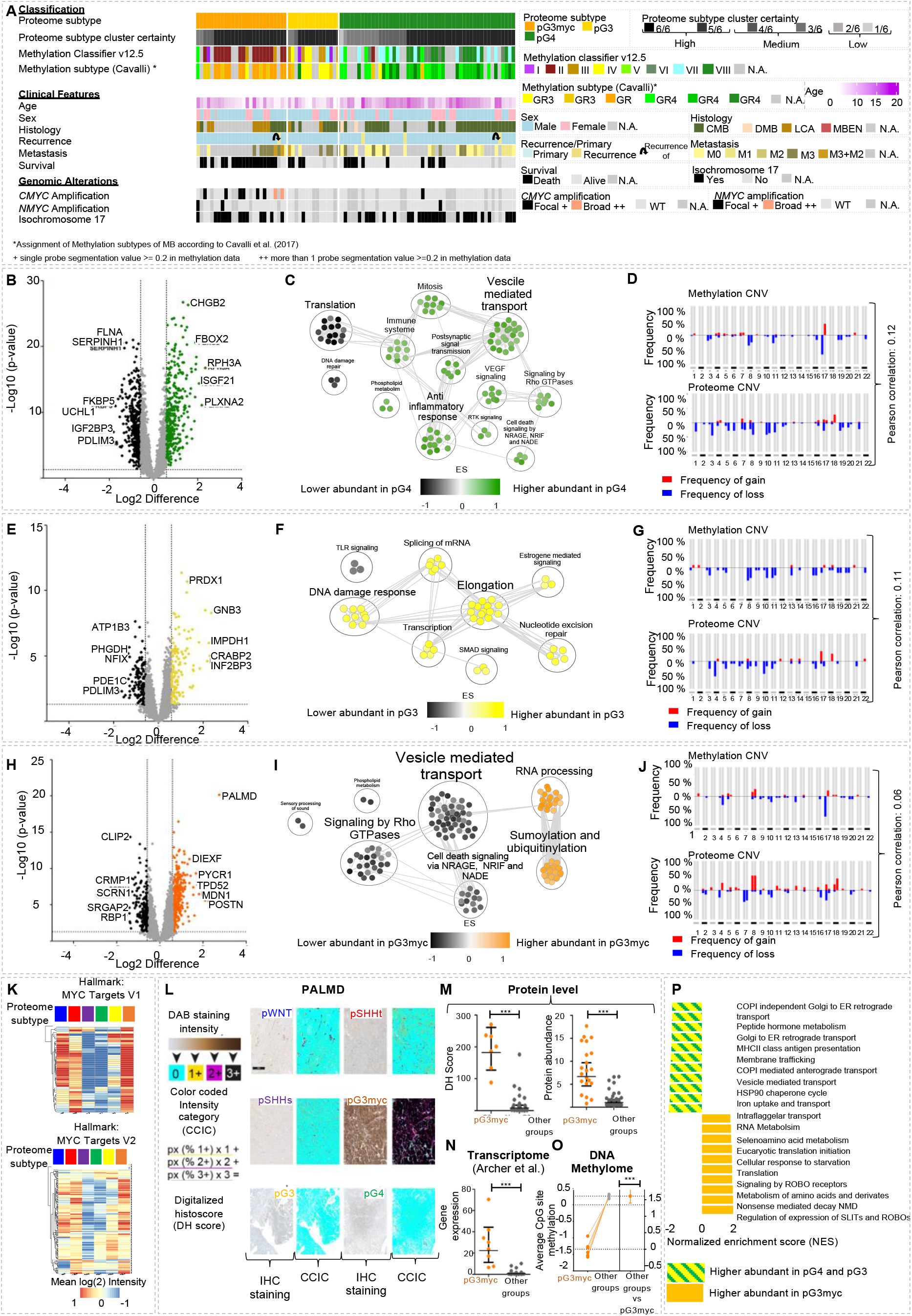
pGroup3-Myc tumors display an enhanced MYC target protein profile and can be identified by Palmdelphin (PALMD) staining. **(A)** Histological, molecular, and clinical characteristics of the MB subtypes pG3myc (n=26), pG3 (n=15) and pG4 (n=40)**. (B)** Volcano plot showing differentially abundant proteins when comparing pG4 tumors to all other proteome subtypes (p-value<0.05; Foldchange difference > 1.5). (**C)** Gene set overlap dependent MCL clustering of enriched gene sets, comparing pG4 to all other subtypes in GSEA. **(D)** CNV plots of pG4 MBs (n=40) calculated from either DNA methylation or proteome data with correlation between both omic types (r=0.12). **(E)** Volcano plot showing significantly differentially abundant proteins when comparing pG3 tumors to all other proteome subtypes (p-value<0.05; Foldchange difference > 1.5). **(F)** Gene set overlap dependent MCL clustering of enriched gene sets, comparing pG3 to all other subtypes in GSEA. **(G)** CNV plots of pG3 MB (n=11) calculated from either DNA methylation or proteome data with correlation between both omic types (r=0.11). **(H)** Volcano plot showing differentially abundant proteins when comparing pG3myc MB to all other proteome subtypes. Palmdelphin (PALMD) was identified as significantly and highly abundant in pG3myc tumors (p-value<0.05; Foldchange difference > 1.5). **(I)** Gene set overlap dependent MCL clustering of enriched gene sets, comparing pG3myc to all other subtypes in GSEA. **(J)** CNV plots of pG3myc MB (n=20) calculated from either DNA methylation or proteome data with correlation between both omic types (r=0.06). **(K)** Heatmaps showing mean protein abundance in MB subtypes for hallmark gene sets MYC Targets V1 and MYC Targets V2. **(L)** Scheme and representative images of digitally supported immunostaining intensity quantification of PALMD immunostainings in MB. Quantified pixels of different staining intensities were used to calculate a digital Histo-score (DHS), **(M)** Significantly enhanced digital histoscore for PALMD in pG3myc MB (n=7) compared to all other MB subtypes (n=22, p<0.0001). **(N)** Protein abundance for PALMD in pG3myc MB (n=21) compared to all other MB subtypes (n=84, p<0.0001). **(O)** PALMD gene expression in pGroup3myc MBs (n=6) compared to all other MB subtypes (n=30, p<0.0001, data extracted from Archer et al. 2018). **(P)** Average DNA methylation at CpG sites of the PALMD gene (Mean M-values of n=6 CpG sites shown). pGroup3myc MBs show significant lower levels of methylation (p<0.0001). **(Q)** Gene set enrichment analyses (GSEA) showing the top 10 up or downregulated pathways comparing pG3myc MB to pG3/4 MB (p<0.01, FDR<0.25). NES=normalized enrichment score.

### Figure 5

Focusing on non-WNT/ non-SHH MB, we found three different proteome subtypes, which we termed pG3, pG3myc and pG4 (Figure 5A). pG4 exclusively included patients assigned to the main molecular subgroup G4, whereas pG3myc was dominated by G3 patients. pG3 included fractions of both molecular subgroups (Figure 2D). Most pG3 and pG4 patients showed a classic histology (CBM), while pG3myc was dominated by large cell anaplastic (LCA) tumors. LCA histology and *MYC* amplification are used as markers for high-risk tumor stratification in non-WNT/ non-SHH MBs^37^. *MYC* amplifications were predominantly detected in pG3myc tumors, but not all pG3myc classified cases were classified as *MYC* amplified. *NMYC* amplification was not restricted to the pG3myc subtype. With respect to metastatic status, most pG3myc tumors were classified as M3, while pG3 and pG4 were dominated by M0 and M1 patients. In concordance with these results, a broad fraction of pG3myc cases were assigned to the methylation subtype II (16/20 cases, 80%)^28,38^ or group G3 δ^5^ (13/20 cases, 65 %, Figure 5A). Conversely, 94.12 % of subtype II cases fell into the pG3myc group. pG3 MBs were mainly composed of methylation subtypes 1, 3 and 4 (10/11 cases, 90.90 %). Finally, pG4 tumors commonly showed the methylation subtypes 5, 6, 7 and 8 (35/37 cases, 94.59 %). All groups showed a low overall correlation between calculated proteome CNV and DNA methylation CNV data (Figure 5D, G, J, Supplementary Table 5j-l). For pG4, we found 167 differentially abundant proteins, including CHGB2, FBOX2, RPH3A, ISGF21 and PLXNA (Figure 5B, Supplementary Table 5c). GSEA revealed an overrepresentation of immune system and synapsis associated processes (FDR< 0.25; p-value <0.0001). Furthermore, a significant overrepresentation (Benjamini Hochberg FDR < 0.01) of VEGF signaling, cell death signaling by NRAGE, NRIF and NADE and phospholipid metabolism was detected (Figure 5C; Supplementary Table 5h, i).

For pG3, 92 differentially abundant proteins included PRDX1, GNB3, IMPDH1, CRAPBP2 and INF2BP3(Figure 5E, Supplementary Table 5b). GSEA identified an enrichment of transcription/translation related processes and DNA repair associated terms (FDR< 0.25; p-value <0.0001). Additionally, estrogen mediated signaling associated genes were overrepresented in pG3 whereas TLR signaling was significantly underrepresented (Figure 5F; Supplementary Table 5f, g).

89 differential abundant proteins were detected inpG3myc MBs, including DIEXF, MDN1, POSTN and PALMD. TPD52, previously proposed as a potential biomarker for high-risk non-WNT/ non-SHH MB^8^, was three times higher abundant in pG3myc (Figure 5H; Supplementary Table 5a). GSEA identified an enrichment for RNA processing and SUMOylation and Ubiquitinoylation related proteins (FDR< 0.25; p-value <0.0001), along with significant underrepresentation of proteins involved in synaptic processes, cell death signaling, phospholipid metabolism and sensory processing (Figure 5I; Supplementary Table 5d, e). In addition, Hallmark gene set-based GSEA revealed a significant enrichment of MYC target proteins in pG3myc (FDR< 0.25; p-value <0.0001, Figure 5K). High abundance levels of Palmdelphin (PALMD) (fold change difference: 6.5, Figure 5 H), were confirmed in pG3myc MBs using digitally supported quantification of PALMD immunostaining (Figure 5L, M). In addition, a significantly higher *PALMD* mRNA expression was detected in reanalyzed data of pG3myc MBs compared to all other MB subtypes (Figure 5N)^18^. CpG sites of the *PALMD* gene showed significant lower levels of methylation in pG3myc MBs and high mRNA expression (Figure 5N). High mRNA expression and low CpG site methylation were associated with poor survival in MB (Figure 5O, Supplementary Figure 7A-D). GSEA focusing on the signaling differences between high-risk pG3myc and other non-WNT/ non-SHH MBs (FDR< 0.25; p-value <0.0001) revealed an enrichment of signaling by ROBO receptors whereas and an underrepresentation of proteins involved in MHCII class antigen presentation or COPI independent Golgi to ER transport (Figure 5P, Supplementary Table 5m-n). In summary, pGmyc MBs were characterized by an enrichment of MYC target proteins, predominantly harbored *MYC* amplifications, and are detectable using PALMD immunohistochemistry.

WNT MB (Figure 6A) displayed 176 differentially abundant proteins including TNC, P4HA2, PTBP3, CABP2 and PMG5 (Figure 6B). TNC showed the highest protein foldchange difference (14.7 foldchange) and its mRNA has been described to be highly expressed in WNT MB^39^ (Figure 6B, Supplementary Table 6a). A significant higher intensity of TNC in pWNT MB was confirmed using digitally supported immunostaining quantification (Figure 6C, D). Using a publicly available dataset^5^, a higher expression of *TNC* in WNT MB was confirmed at the transcriptome level (Figure 6E). CpG sites of the *TNC* gene, measured with the Illumina 850K array, showed no significant difference of methylation (pWNT versus other subtypes (Figure 6F, Supplementary Figure 7). In line with this none of TNC’s CpG sites were identified as potential biomarkers for WNT MB (Figure 3C). GSEA revealed, an enrichment of extracellular matrix proteins and N-glycan biogenesis and transport (FDR< 0.25; p-value <0.0001, Figure 6G, H). Metabolic processes, synaptic signaling and proteins of the TriC/CCT complex were underrepresented (Figure 6G, Supplementary Table 6b, c). The TriC/CCT complex has previously been reported to be associated with vincristine resistance^40^. Typical chemotherapy regimens for MB consist of cisplatin/carboplatin-vincristine-cyclophosphamide combinations^41^. Of note, pWNT MBs showed the lowest abundance of TriC/CCt proteins, whereas pG3myc MBs displayed the highest amount. No such correlation was observed at DNA-methylation or transcriptome level (Figure 6I, Supplementary Table 6e). A high overall correlation between copy number plots extracted from proteome and DNA methylation data was observed for pWNT compared to all other subtypes (Figure 6J, Supplementary Table 6d), being in line with a general increased overall correlation of proteome and DNA methylome data (Figure 3). Taken together, compared to other subtypes, pWNT MB showed the highest correlation of proteome and DNA methylome data and were characterized by low TriC/CCt proteins, as well as high abundance of TNC.

**Figure 6:**
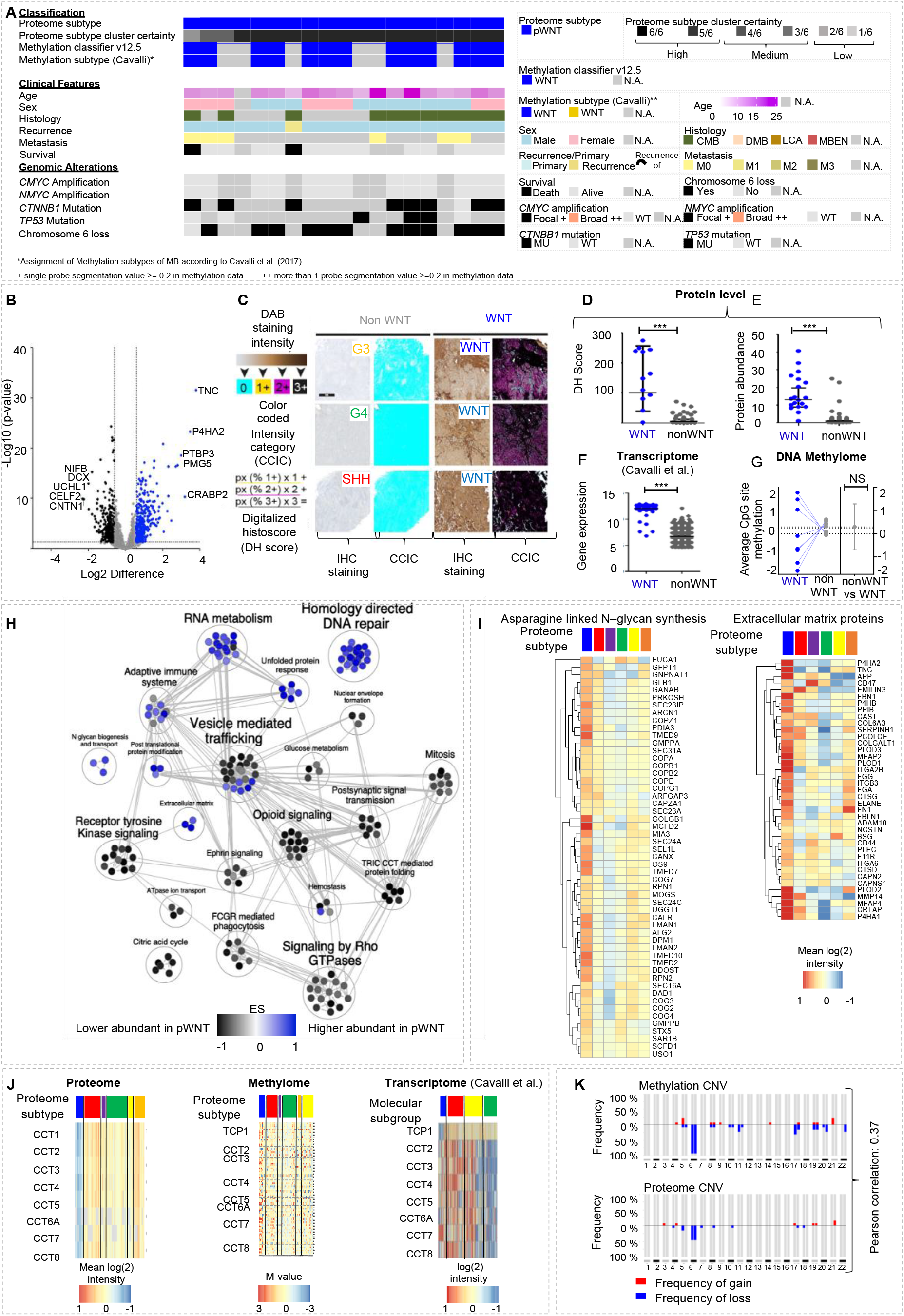
pWNT MB show alterations of the multiprotein complex TriC/CCT and can be identified by Tenascin C (TNC) staining. ***(A)** Histological, molecular, and clinical characteristics of the pWNT MB subtype (n=19). **(B)** Volcano plot showing differentially abundant proteins when comparing pWNT tumors to all other proteome subtypes (p-value<0.05; Foldchange difference > 1.5). TNC was identified as highly abundant in pWNT MB. **(C)** Scheme and representative images of digital quantification of TNC immunostainings in MB. **(D)** Significantly enhanced DHS for TNC in pWNT MB (n=12) compared to all other MB subtypes (n=27, p<0.0001). **(E)** Protein abundance for TNC in pWNT MBs (n=19) compared to all other MB subtypes (n=143, p=<0.0001). **(F)** TNC gene expression in WNT MBs and other MB subtypes in a published dataset of MB^5^. **(G)** Average DNA methylation at CpG sites of the TNC gene (mean value for n=8 CpG sites shown). **(H)** Gene set overlap dependent MCL clustering of enriched gene sets, comparing pWNT to all other subtypes in GSEA **(I)** Heatmaps showing mean protein abundance in MB subtypes for hallmark genesets specifically enriched in pWNT MB (NES_Glycan_=2.2, p_Glycan_=<0.001; NES_EMP_=1.7, p_EMP_=0.02). **(J)** Heatmap showing protein abundancies for components of the tailless complex polypeptide 1 ring complex/ Chaperonin containing tailless complex polypeptide 1 (TriC/CCT) in MB subtypes (left). DNA methylation at CpG sites of TriC/CCT genes in proteome MB subtypes (M-values, middle). Gene expression of TriC*/CCT components *in WNT MBs and other MB subtypes in a published dataset of MB^5^(right). **(K)** CNV plots of pWNT MBs (n=13) calculated from either DNA methylation or proteome data (Pearson, r=0.37)*.

### Figure 7

**Figure 7:**
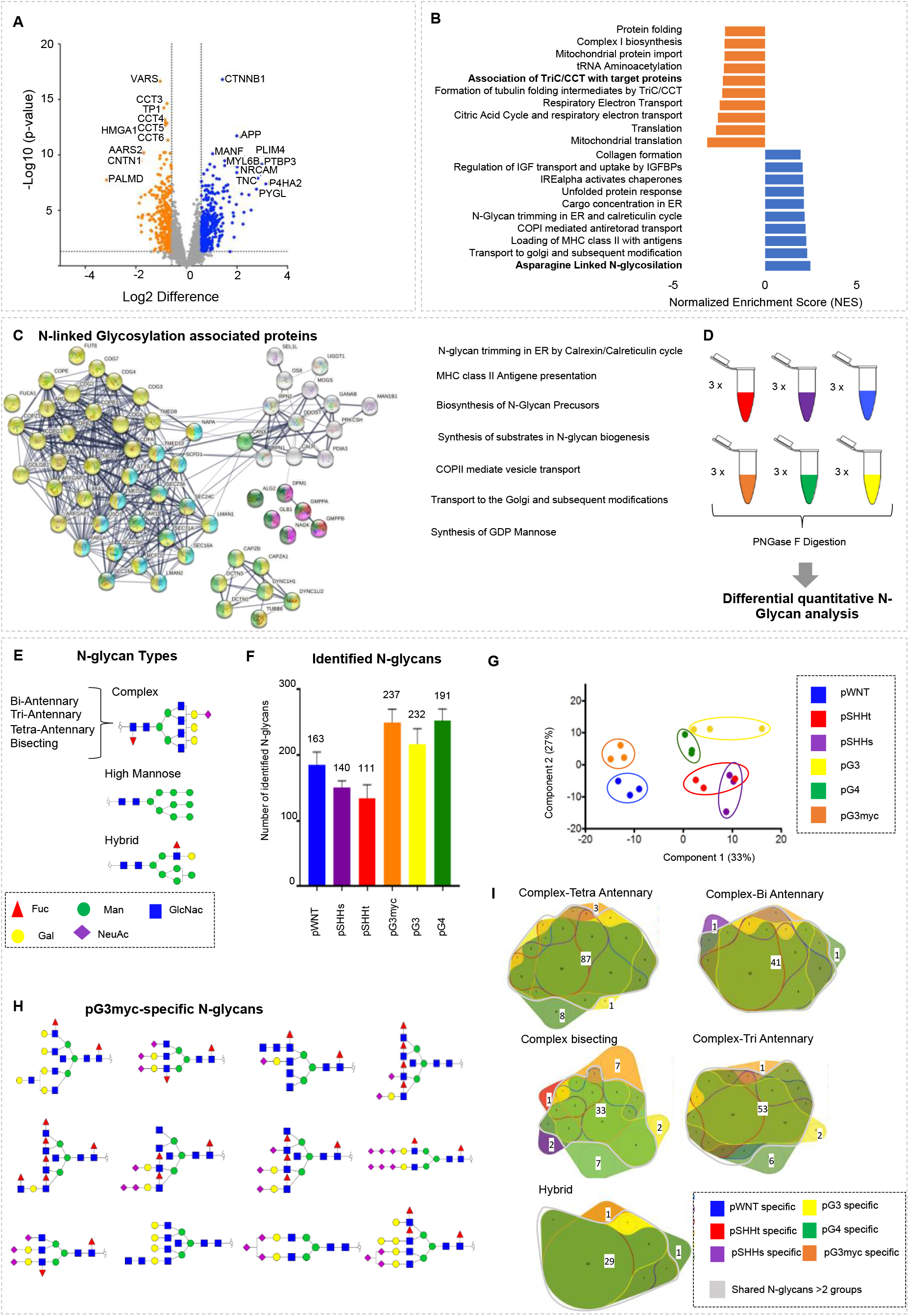
N-glycan analysis reveals significant differences across N-glycan profiles of proteomic MB subtypes. **(A)** Volcano plot depicting differentially abundant proteins when comparing pWNT (n=19) to pG3myc (n=26) MB (p-value<0.05; Foldchange difference > 1.5). **(B)** GSEA showing the top 10 up or downregulated pathways comparing pG3myc MB to pWNT (p<0.05, FDR<0.25). **(C)** STRING network analyses of significant differentially abundant proteins involved in N-linked glycosylation. **(D)** Scheme of N-glycan analyses. **(E)** Schematic visualization of N-glycan types. **(F)** Identified glycans per proteome MB subtype (Error bars represent mean values with standard deviation (SD). The number of commonly identified N-glycans in all samples for each subtype is indicated). **(G)** NIPALS PCA, based on N-glycan abundances, illustrating the separation of proteome MB subtypes at the N-glycan level. **(H)** 2D Structure visualization for pG3myc-specific N-glycans. GlcNAc=N-Acetylglucosamine; Gal=Galactose; Fuc=Fucose; ManNAc=N-Acetylmannosamine; Neu5AC=N-Acetylneuraminic acid. **(I)** Venn Diagram, comparing the identified hybrid-Type and complex N-glycans between proteome subtypes.

The highest similarity of proteome profiles was observed for the pG3myc subtype, associated with high-risk features and the pWNT subtype-associated with relatively good overall survival (Figure 2G). Both subtypes showed a main “transcriptional/translational” profile (Figure 2 G, H) and a high abundance of MYC target proteins (Figure 5K). We therefore asked, what molecular changes could impact on such diverse clinical behavior. In total, 188 proteins were differentially abundant between pG3myc and pWNTMBs including TriC/CCt proteins (CCT2, TCP1, CCT4, CCT2, CCT5, CCT6, CCT8) and the established WNT MB marker β-Catenin (*CTNNB1*)^41^ (Figure 7A, Supplementary Table 7a). GSEA (FDR< 0.25; p-value <0.0001) revealed, that the top 10 enriched gene sets for pG3myc were protein folding, translation, and metabolic processes (Supplementary Table 7b-c). Additionally, asparagine linked N-glycosylation associated factors showed a higher abundance in pWNT tumors compared to pG3myc MBs (Figure 7B). We further focused on this PTM and found that proteins involved in all aspects of N-glycosylation (synthesis, processing, transport and antigen presentation via MHC class II) were overrepresented in pWNT tumors (Figure 7C).

A differential, quantitative analysis of N-glycans revealed differential N-glycosylation patterns across proteomic MB subtypes (Figure 7D-I). In total 302 different N-Glycan species were identified (high mannose: n = 11; complex-bisecting: n=51, complex-bi-antennary: n = 46, complex-tri-antennary: n = 64, complex-tetra-antennary: n=99, hybrid: n=31, Figure 7 E-I; Supplementary Table 7d). For non-WNT/ non-SHH MB a significantly higher number of N-glycans were identified in comparison to pWNT, pSHHs and pSHHt (Figure 7F, Supplementary Table 7d). At the quantitative level, proteome MB subtypes were also reflected based on their N-glycan profiles, with SHH subtypes being most similar to each other (Figure 7G, Supplementary Figure 8A). 92 N-glycans were differentially abundant between the proteome MB types (Supplementary Figure 8B, Supplementary Table 7e). Comparing identified N-glycan types between proteome MB subtypes, we identified the highest number of exclusive N-glycans for the subtypes pG3myc *and pG4* (n_pG3nyc_ = 22, n_pG4_ = 12). In the pG3myc subtype, several complex-bisecting N-glycans were exclusively identified (Figure 7H, I). In pG4, different complex-bisecting or complex-tetra-antennary N-glycans were found exclusively (Figure 7I). Frequently described key factors in tumors are the upregulation of cancer associated sialynated N-glycans as well as aberrant fucosylation^42^. Focusing on these aspects across MB groups, a significantly higher proportion of sialynated N-glycans was found in non-WNT/ non-SHH tumors (pWNT (51.9% (n = 84)), pSHHs (50.1% (n = 71)), pSHHt (49.5% (n = 55)), pG3myc (59.7% (n = 139)), pG4 (62% (n = 144)), pG3 (60.7% (n = 116))). A significantly lower proportion of fucosylated N-glycans was detected in pSHHt, compared to all other subtypes (66.7 % (n = 74)) versus 72.1 - 80% (n = 101-174, range of the other MB subtypes). In summary, the N-glycome was significantly different among proteome subtypes, forming a basis for new biomarker discovery, based on N-glycan profiles of brain cancers.

## Discussion

DNA methylome and transcriptome analysis, are used for diagnostics in neuro-oncology for the classification of brain tumor types^1^. In contrast to these omics-types, the proteome, and its modifications better reflect the pharmacologically addressable phenotype and may disclose phenotypically relevant processes, potential biomarkers, and therapeutic targets.

In this study, we show the proteome analysis of MB from FFPE material, that maintains chemical rigidity under cheap storage conditions ^43^. We found that FFPE material is suitable for the differential proteome analysis of MB, enabling a clear differentiation between the 4 main molecular subtypes, that was previously described for smaller cohorts of FF tissue^18,19^. In line with previous results^44^, we found that the sample age (spanning a period of 50 years) did not impact on the data quality, making FFPE tissue highly suitable for large-scale analysis of rare diseases.

Using the HarmonizR strategy for the integration of independently generated proteome datasets^27^, we established an integrated dataset of 167 MBs. We identified six different proteome MB subtypes, in line with previously published results, obtained from smaller fractions of the analyzed cases^18^. WNT tumors formed a common cluster. For hedgehog driven MB cases two independent clusters were formed (pSHHt, pSHHs), while non-WNT/ non-SHH tumors diverged into three subtypes (pG3, pG4, pG3myc). Interestingly, two overriding molecular patterns were observed across MB subtypes, indicating that MB either follow a transcriptional/replicative (pWNT, pSHHt, pG3myc) or synaptic/ immunological (pG4, pSHHs, pG3) profile.

### Integration of proteome and DNA methylation data

Proteome subtypes partly overlapped with previously defined DNA methylation subtypes^4–6,31,28^. Specifically, pG3myc tumors were mainly composed of methylation subtype II. pG3 tumors comprised mainly methylation subtypes 1, 3 and 4 and pG4 tumors showed methylation subtypes 5, 6, 7 and 8. Finally, the DNA methylation subtypes SHH3 and SHH4 were exclusively found in pSHHt MBs. Only 30% of marker proteins showed a significant correlation with their respective gene’s CpG sites. In general, a low correlation between proteome and methylome was found in MB, in line with results of previous studies on other tumor entities^45,46^. Poor correlations might also be attributed to the 850K array design, since it mostly assesses promoter methylation sites whereas CpG sites correlating well with gene expression may locate further away from transcriptional start sites^47^. Of note, correlation levels of data modalities were not evenly distributed among subtypes. Especially proteins detected in pWNT tumors, showed a high correlation with their respective gene’s CpG sites (38.9% of proteins). In addition, the commonly detected loss of chromosome 6^48^ was also reflected in proteome data when mapping protein abundances to chromosomal arms. This indicates that molecular alterations may be more conserved for WNT MBs, whereas DNA-based methylation differences do not always result in an effective change in protein abundance, highlighting the importance of proteome analysis.

### Proteome subtypes of SHH-MB

SHH MBs divided into two proteome subtypes namely, pSHHs and pSHHt, confirming previous results from a smaller MB dataset^18^. The pSHHs tumors reflect the SHHb subgroup defined by Archer et al showing an enrichment of synaptic pathways^18^. We found that these tumors are characterized by high representation of the citric acid (CTA) cycle and respiratory electron transport, pointing at a distinct metabolic profile of pSHHs MB. *TP*53-mutated SHH are stratified as high-risk SHH MB^36^. Based on the proteins considered here, *TP*53-mutated cases did not form a distinguishable cluster. However, among others CHD6, DNAJB2 and NNMT, known to be associated with aberrant *TP*53 expression and high tumor progression ^49–51^, showed a differential abundance comparing *TP*53-mutated to *TP*53-wildtype cases. Further, CHD6 is suggested as a potential anti-cancer target for tumors with DNA-damage repair associated processes^50^. Mutations within the largest subunit of the elongator complex (*ELP1*) have lately been described in SHH MB^33^. These mutations were found mutually exclusive with *TP53* mutations and *ELP1* mutated SHH MBs were characterized by translational deregulation with upregulation of factors involved in transcription and translation^33^. A reanalysis of published proteome data from *ELP1* mutated SHH MB cases indeed revealed that all cases were attributed to the pSHHt MB subtype (Supplementary Table 4k)^33^. As a limitation, the *ELP1* status of the SHH MB cases in our cohort was only known for n = 3 pSHHs and n = 10 pSHHt tumors (all wildtype). However, all SHH MBs with methylation subtype 3 - associated with *ELP1* mutations - fell into pSHHt ^28,33^.

### pWNT

The current standard treatment for MB is surgical removal of the tumor followed by craniospinal irradiation and combinational chemotherapy. These approaches cause severe late effects, mostly neuro-cognitive and neuroendocrine sequelae. Due to their high responsiveness to therapy, WNT-type MBs are currently being evaluated for therapy de-escalation, making their clear identification indespensable^52^. On a molecular level the identification of *CTNNB1* mutations, or chromosome 6 deletion (monosomy 6) are common markers for the identification of WNT-type MB. Immunohistochemistry is used to detect nuclear ß-Catenin staining in tumor cells. However, nuclear staining can be weak and only a subset of cell nuclei is usually stained^53,54^. Here, Tenascin C (TNC) was found elevated in pWNT MBs from proteome and mRNA data, whereas no significantly altered DNA methylation at CpG sites of *TNC* was seen.. *TNC* is a highly glycosylated extracellular matrix (ECM) protein, promoting or inhibiting proliferation and migration in cancer, depending on the present splice variant^55^, which will be a field of further study. Besides TNC, a general enrichment of ECM proteins was detected in pWNT MBs. While the ECM has not been investigated in-depth in WNT MB, ECM components have been described to predict patient outcomes in MB^56^. ECM degradation was found as a hallmark of tumor invasion, metastasis development and overall bad prognosis^57^. WNT pathway activation dependent disruption of the blood-brain barrier (BBB)^57^, was described to permit accumulation of high levels of intra-tumoral chemotherapy in WNT tumors, resulting in a robust therapeutic response. TNC could be another contributor to this phenotype, as high TNC levels contribute to BBB disruption^57,58^. Furthermore, other BBB contributors, such as *EPLIN1*, DSP and S100A4 were found differential in pWNT (Supplementary Table 7a).

### Non-WNT/ non-SHH MB

In line with previous results, we found three proteome subtypes of non-WNT/ non-SHH MBs^18^. pG4 (predominantly consistent of main molecular subtype G4 tumors), followed the synaptic program. These findings go in line with the literature, as synaptic signatures for G4 tumors, have been previously described^5,18^. In pG4 MBs, we also detected a higher abundance of VEGF signaling-related proteins, previously described in the context of tumor angiogenesis. VEGF signaling can be targeted in MB using Bevacicumab or Mebendazole ^59,60^ and hence might be especially be beneficial for pG4 patients.

pG3 MBs (composed of both G3 and G4 tumors) showed the lowest cluster certainty and inherited the characteristics of both pG3myc and pG4 tumors.

pG3myc tumors, showed a reduced survival rate and high-risk features, such as LCA histology and solid metastasis. Group 3 MB with *MYC* amplification have been shown to be highly aggressive and exhibit a bad prognosis^61,62^. In our cohort, more than half of the patients showed a *CMYC* amplification, while all samples showed an upregulation of *CMYC* target genes, supporting the hypothesis that besides *CMYC* amplification, changes in its phosphorylation status result in a CMYC-*driven* high-risk proteome G3 subtype^18^. For this group, we identified, a significant enrichment of signaling by ROBO receptors, amino acid metabolism, RNA metabolism, and nonsense-mediated RNA decay and translation. As potential protein biomarkers *DIEXF, MDN1, POSTN, TPD52 and PALMD* showed a higher abundance in pG3myc tumors. *TPD52* has recently been suggested as an immunohistochemistry (IHC) marker for high-risk non-WNT/ non-SHH patients^8^. PALMD showed the highest elevation in pG3myc MB in our cohort and was established as a suitable IHC marker for the identification of pG3myc MB. However, for biomarker validation further prospective trials are needed to evaluate its significance for stratification of high-risk non-WNT/ non-SHH patients.

### pG3myc and WNT

High-risk pG3myc MBs showed a high resemblance to pWNT tumors, that are associated with a favorable outcome. Comparing both groups, proteins associated with Tric/CCT complex were elevated in pG3myc MBs. A high abundance of CCT complex proteins has previously been linked to worse prognosis in cancer and was identified as a predominate driver of Vincaalcaloid resistance, including Vincristine, which is among the most frequently used drugs for all MB subtypes^63^.The general low abundance of CCT/TriC proteins in pWNT MB could therefore be a BBB-phenotype independent explanation for the relatively high response to chemotherapy^64^. The usage of CT20p, an amphipathic CCT inhibitor peptide, was recently described as a promising strategy for the treatment of high-risk tumors with high CCT abundance^65,66^. Based on our data, the approach should be further investigated as a potential strategy to enhance Vincristine-mediated cytotoxicity in high-risk non-WNT/ non-SHH MBs.

We further identified increased Asparagine-linked-N-glycosylation as a hallmark of WNT Medulloblastoma. While aberrant N-glycosylation patterns have been described for brain cancer, especially focusing on sialylation and fucosylation^67^, a global analysis of N-glycans has not been performed yet. It is reasonable that certain glycosylation patterns can be used as biomarkers for the progression of diseases^22^. In addition, aberrant N-glycan structures in cancer could be targeted by immunotherapy and provide new therapeutic strategies, especially for high-risk tumors that are not sensitive to classical treatment^68,69^. As an example, chimeric-antigen-rceptor (CAR)-modified T cells, that can be specifically directed against tumor-associated carbohydrate antigens (TACAs) are rapidly evolving^70^. Differential, quantitative N-glycan analysis, identified a total of 303 N-glycans in MB. Based on the quantitative distribution of commonly found N-glycans in all samples, the proteome subgroups were reflected. pSHHt and pSHHs MBs were most similar. This could be related to the dominant SHH activation in these groups, knowingly having a severe impact on N-glycosylation^71^. 12 structures were identified only in high-risk pG3myc patients. Most of these structures are complex bisecting N-glycans, that are known to be associated with cell growth control and tumor progression^22,71^ and might be related to the unfavorable outcome for pG3myc patients. pG3myc-specific N-glycans do not appear healthy brain cells, whose N-glycome is characterized by dismissed N-glycan complexity, lack of complex N-glycans and truncated structures^72^ and might serve as suitable immunotherapy targets for high-risk patients.

For pG4 patients, the highest amount of salivated N-Glycans was found, further supporting the immunological profile of pG4 MBs, observed at the proteome level^73^.

Taken together, the integration of MB proteome, DNA-methylome and N-glycome revealed new insights into MB phenotypes, potential new biomarkers and therapeutic targets for MB such as the usage of TriC/CCT-inhibitors and chimeric-antigen-receptor-modified T-cells to target tumor-specific carbohydrates for high-risk MB patients.

## Author contributions

H.V., S.G. and J.N. wrote and reviewed the manuscript. J.N. planned and designed the study. H.V and S.G. conducted experiments. H.V, S.S., B.P., T.M., H.S., C.K. and Y.G. analyzed proteome and glycosylation data. U.S. and S.G analyzed methylation data. S.G. and Y.S. integrated proteome and methylome data. M.D. performed digitally supported quantification if IHC. S.P., S.R., M.M., MM. D., A.K., C.H., J.W., and F.L-S. analyzed and interpreted histological, molecular and clinical data. All authors reviewed the manuscript and approved its final version.

## Acknowledgements

We thank Tasja Lempertz, Carolina Janko, Ulrike Rumpf and Karin Gehlken for skillful and kind support. JN is funded by the Deutsche Forschungsgemeinschaft (DFG, Emmy Noether program).

## Inclusion and Diversity

We support inclusive, diverse, and equitable conduct of research.

## Methods

### Subject Details

#### In house patient samples

FFPE Medulloblastoma samples of tumors within the years 1976-2021 were obtained from tissue archives from various neuropathology units in Germany including cases that had been collected within the HIT-MED study cohort. All investigations were performed in accordance with local and national ethical rules of patient’s material and have, therefore, been performed in accordance with the ethical standards laid down in an appropriate version of the 1964 Declaration of Helsinki. All samples underwent anonymization. Tumor samples were fixed in 4 % paraformaldehyde, dehydrated, embedded in paraffin, and sectioned at 10 μm for microdissection using standard laboratory protocols. For further information on clinical details of samples, please refer to Supplementary Table 1c.

#### Medulloblastoma cell lines

The human Medulloblastoma cell lines DAOY (Ca#HTB-186) and D283med (Ca#HTB-185) were obtained from ATCC, Manassas, VA, USA. UW473 was kindly provided by Michael Bobola. All lines were used as Standards for TMT batches. Cells were cultivated in DMEM (Dulbecco’s Modified Eagle Medium, PAN-Biontech) supplemented with 10 % FCS at 37°C, 5 % CO2.

#### Publicly available datasets

For the data integration and harmonization of in-house and publicly available DNA Methylation data the following datasets were used: Archer et al. (2018)^18^: 42 FF MB samples, accessible as a subset of European Genome-phenome Archive ID: EGAS00001001953. Forget et al. (2018)^19^: 38 FF MB samples, accessible via Gene Expression Omnibus (GSE104728). For the analysis of RNA Expression data, processed and normalized data from the following datasets were used: Cavalli et al. (2017)^18^: 763 MB samples, accessible via Gene Expression Omnibus (GPL22286)^5^. For the data integration and harmonization of in-house and publicly available proteome data, the following datasets were included: Archer et al. (2018)^18^: 45 FF MB samples, available via the MassIVE online repository (MSV000082644, Tandem Mass Tag- (TMT) label-based protein quantification); Forget et al. (2018)^19^: 39 FF MB samples,, available via the PRIDE archive (PXD006607, stable isotope labling by amino acids in cell culture- (SILAC) label-based protein quantification); Petralia et al. (2021)^17^, 23 FF MB samples,, available through the Clinical Proteomic Tumor Analysis Consortium Data Portal (https://cptac-data-portal.georgetown.edu/cptacPublic/) and the Proteomics Data Commons (https://pdc.cancer.gov/pdc/, Tandem Mass Tag- (TMT) label-based protein quantification). For validation of determined proteome subtypes, as well as the investigation of the proteome profile of ELP1 mutated SHH MB, a dataset published by Waszak et al. (2020)^33^ was used (23 FF MB samples, available via the PRIDE archive (PXD016832, Data independent acquisition label free protein quantification).

### Sample preparation and data acquisition

#### DNA methylation profiling

DNA methylation data was generated from FFPE tissue samples. DNA was isolated using the ReliaPrep™ FFPE gDNA Miniprep system (Promega) following the manufacturer’s instructions. 100-500 ng DNA was used for bisulfite conversation using the EZ DNA Methylation Kit (Zymo Research). Then the DNA Clean & Concentrator-5 (Zymo Research) and the Infinium HD FFPE DNA Restore Kit (Illumina) were applied. Infinium BeadChip array (EPIC) using manufacturer’s instructions were then used to quantify the methylation status of CpG sites on an iScan (Illumina, San Diego, USA). Additionally, previously published data measured on Infinium Human Methylation 450 BeadChip array (450K) were included from EGAS0001001953^74^, from GSE104728^19^, and GSE130051^75^.

#### Proteome profiling

FFPE MB tissue sections were deparaffinized with N-heptane for 10 minutes and centrifuged for 10 minutes at 14,000 g. The supernatant was discarded. Proteins were extracted in 0.1 M triethyl ammonium bicarbonate buffer (TEAB) with 1% sodium deoxycholate. (SDC) at 99°C for 1hour. Sonification was performed for 10 pulses at 30% power, to degrade DNA, using a PowerPac™ HC High-Current power supply (Biorad Laboratories, Hercules, USA)) probe sonicator. For cell lines, proteins were extracted in 0.1M triethyl ammonium bicarbonate buffer (TEAB) with 1% sodium deoxycholate. (SDC) at 99 °C for 5 minutes. Sonification was performed for 6 pulses.

The protein concentration of denatured proteins was determined by the Pierce BCA Protein assay kit (Thermo Fischer Scientific, Waltham, USA), following the manufacturer’s instructions. 60 μg of protein for each tissue lysate and 30 μg protein for each cell lysate were used for tryptic digestion. Disulfide bonds were reduced, using 10mM dithiothreitol (DTT) for 30 minutes at 60 °C. Alkylation was achieved with 20 mM iodoacetamide (IAA) for 30 minutes at 37 °C in the dark. Tryptic digestion was performed at a trypsin: protein ratio of 1: 100 overnight at 37 °C and stopped by adding 1% formic acid (FA). Centrifugation was performed for 10 minutes at 14000g to pellet precipitated SDC. The supernatant was dried in a vacuum concentrator (SpeedVac SC110 Savant, (Thermo Fisher Scientific, Bremen, Germany)) and stored at −80°C until further analysis.

For the main cohort, 50 μg sample per patient and internal reference, TMT-10 plex labeling (Thermo Fischer Scientific, Waltham, USA), was performed, following the manufacturer’s instruction. All 70 patients were run in 8 total TMT 10-plexes. Samples assignment to batches was performed in a semi-randomized manner, according to the four main molecular subtypes. In each batch, 1-2 internal reference samples were included, composed of equal amounts of peptide material from all 70 samples and cell lines. Isobarically labeled peptides were combined and fractionated, using high pH reversed phase chromatography (ProSwiftTM RP-4H, Thermo Fischer Scientific Bremen, Germany) on a HPLC system (Aglient 12000 series, Aglient Technologies, Santa Crara, USA). Separation was performed using buffer A (10 mM ammonium hydrogen carbonate (NH_4_HCO_3_) in in H2O) and buffer B (10mM NH_4_HCO_3_ in ACN) within a 25-minute gradient, linearly increasing from 3-35% buffer B at a flow rate of 200 nl/min. In total, 13 fractions were collected for each batch, dried in a vacuum concentrator (SpeedVac SC110 Savant, (Thermo Fisher Scientific, Bremen, Germany)), resuspended in 0.1 % FA to a final concentration of 1mg/ml and subjected to high pH liquid chromatography coupled mass spectrometry (LC-MS). All LC-MS measurements were performed on a UPLC system (Dionex Ultimate 3000, Thermo Fisher Scientific, Bremen, Germany, trapping column: Acclaim PepMap 100 C18 trap ((100 μm x 2 cm, 100 Å pore size,5 μm particle size); Thermo Fisher Scientific, Bremen, Germany), analytical column: Acclaim PepMap 100 C18 analytical column ((75 μm x 50 cm, 100 Å pore size, 2 μm particle size); Thermo Fisher Scientific, Bremen, Germany)), coupled to an quadrupole-orbitrap-iontrap mass spectrometer (Orbitrap Fusion, Thermo Fisher Scientific, Bremen, Germany). Separation was performed using buffer A (0.1% FA in H20) and buffer B (0.1% FA in H20) within a 60-minute gradient, linearly increasing from 2-30% buffer B at a flow rate of 300nl/min. Eluting peptides were analyzed, using a DDA based MS3 method with synchronous precursor selection (SPS), as described by McAlister et al.^76^. For further details on protein extraction, tryptic digestion, and LC-MS/MS setups, please refer to the PRIDE archive (PXD039319).

#### N-Glycan profiling

100 ug of protein for 18 samples was denatured, reduced, and alkylated as described above. Samples was concentrated by 3 kDa Amicon Ultra centrifugal filters (Merck Millipore, R0NB30416) with 100 mM NH_4_HCO_3_ to exchange the buffer and retain globular particles above 3 kDa. Thirty units of PNGase F were added to each sample and incubated in a 37 °C Thermomixer for 24 h. After PNGase F digestion, purified N-glycans were eluted by Sep-Pak C18 cartridges (Water, WAT023590) with 5% acetic acid and dried in a speed vacuum. 40 μl of 10 μg/μL (w/v) ammonium borane solution was added to purified N-glycan samples and incubated in a 60 °C Thermomixer for one h. After incubation, the borane-ammonia complex was removed by the repeated evaporation of 400 μl of methanol using a speed vacuum. The reducing N-glycan sample was permethylated by optimized solid-phase permethylation as describe by Guan et al.(2020)^77^(76). Purified and reduced N-glycan samples were dissolved in 110 μl of DMSO/water (100:10) solution, and then 70 μl of methyl iodide was added. Redissolved samples were transferred to a tube which contained 200 mg sodium hydroxide beads and incubated in a Thermomixer for 10 min by 1300 rpm at room temperature. 200 μl of 5% acetic acid was added to each sample to quench the permethylation and eliminate oxidation reactions. Derivatized N-glycan was isolated using 300 μL of chloroform by chloroform-water extraction. Permethylated reducing N-glycans were resuspended in 0.1% FA solution to a final concentration of 2 mg/ml and subjected to LC-MS. For N-glycans, separation was achieved using a linear gradient from 2% to 30% buffer B in 10 min, increased to 70% in 75 min and finally to 95% in 85 min. Ions were detected in automatic gain control (AGC) (target: 2.0e5 ions) with an accumulation time of 120 ms. Induced collision was performed at 35% normalised collision energy and an isolation window of 2 m/z).

### Raw data processing

#### Processing of DNA Methylation Array Data

Idat files generated using the above protocol were processed in R (Version 4.0.5). The files were read in using the minfi package (Version 1.36.0)^78^.Detection P-value was used to identify sample quality and filter out bad quality samples (none were excluded, n=0). Further, probes having bad quality (n=49,091), probes with single nucleotide polymorphism (n = 12,868) and probes present on X and Y chromosomes (n=8,777) were filtered out. After normalization and probe filtering, the m-values log2(M/U) where methylation intensity is denoted by M and unmethylation intensity denoted by U were used for further analysis. Differentially methylated probes/ CpG sites were found using the limma package (Version 3.46.0)^79^, comparing all subtypes using the contrast function and correcting for multiple testing using Benjamini Hochberg (cut-off 5% FDR). Based on this, M-values of 10,000 differentially methylated CpG sites which could cluster subtypes based on biological differences were selected for further analysis. Heatmaps were generated using ComplexHeatmap (Version 2.6.2)^80^ and pheatmap (Version 1.0.12).

#### Processing of Proteome raw data

Obtained raw data from in-house generated and publicly available (Archer et al (2018)^18^, TMT 10-Plex; Petralia et al. (2021)^17^, TMT 11-Plex). TMT-based LC-MS measurements were processed with the Andromeda algorithm, implemented in the MaxQuant software (Max Plank Institute for Biochemistry, Version 1.6.2.10)^81^ and searched against a reviewed human database (downloaded from Uniprot February 2019, 26,659 entries). The Carboxymethylation of cysteine residues was set as a fixed modification. Methionine oxidation, N-terminal protein acetylation and the conversion of glutamine to pyroglutamate were set as variable modifications. Peptides with a minimum length of 6 amino acids and a maximum mass of 6,000 Da were considered. The mass tolerance was set to 10 ppm. The maximum number of allowed missed cleavages in tryptic digestion was two. A false discovery rate (FDR) value threshold <0.01, using a reverted decoy peptide databases approach, was set for peptide identification. Quantification was performed, based on TMT reporter intensities at MS3 level for LC-MS3 in-house data and at MS2 level for LC-MS2 data, acquired by Archer et al.^18^ and Petralia et al.^17^ All studies were searched separately. Fractions for each TMT batch were searched jointly.

For stable isotope labeling by amino acids in cell culture (super-SILAC) data, acquired by Forget at al. (2018)^19^, log2 transformed SILAC ratios were directly obtained from the MassIVE online repository (MSV000082644).

For the external validation the dataset published by Waszak et al. (2020)^33^ was used. The DIA raw data spectra were downloaded from PRIDE and processed using Data Independent Acquisition with Neural Networks (DIA-NN, version 1.8.1) ^82^. The spectra were searched against a peer reviewed human FASTA database (downloaded from UniProt April 2020, 20,365 entries). A spectral library was generated in silico by DIA-NN using the same FASTA database. Smart profiling was enabled for library generation. Methionine oxidation, carboxymethylation of cysteine residues as well as N-terminal methionine excision were set as variable modifications. The maximum number of variable modifications was set to three, the maximum number of missed cleavages was two. The peptide length range was set from 7 to 30. Mass accuracy, MS1 accuracy, and the scan window were optimized by DIA-NN. An FDR < 0.01 was applied at the precursor level - decoys were generated by mutating target precursors’ amino acids adjacent to the peptide termini. Interference removal from fragment elution curves as well as normalization were disabled. Neural network classifier was set to single-pass mode and the fixed-width center of each elution peak was used for quantification.

#### Processing of N-Glycan raw data

N-Glycan raw data were open with Xcalibur Qual Browser (Version No 4.2.28.14). MaxQuant were used for extracting all the detected masses and *m/z* from MS raw data of permethylated reducing N-glycans. Home-made Python-scripts is used to extract and calculate monosaccharide compositions based on the molecular weight of each derivatized N-glycan^83^. The N-glycan structures were identified manually based on full MS, *m/z* and MS2 ion according to Xcalibur and Glycoworkbench 2.1. The monosaccharide compositions, *m/z* and charge were exported into the Skyline software (Version No 21.1.0.278) to calculate the peak area of each N-glycan. Finally, the table, including N-Glycan compositions, the abundance of N-glycan and mass, was input into the Perseus software.

### Data normalization and integration

#### Normalization and integration of DNA Methylation Array Data

Single-sample noob normalization (ssNoob) was performed since we combined samples from different arrays (EPIC and 450K). In single sample noob normalization, there is no need for reference sample-based dye-correction. The detailed method development has been mentioned^84,85^. Raw signal intensities for EPIC and 450K files were read individually. Since ~ 93% of the loci of 450K array are also present on EPIC array, they can be combined using minfi’s combineArrays (). After combining the two arrays they can be output as a virtual array. In this study, 450K array was the output virtual array since a greater number of samples were measured on 450K.

#### Normalization and integration of Proteome data

Prior to data integration, protein abundances were handled separately for each dataset. TMT reporter intensities were log2 transformed and median normalized across columns. Technical variances between TMT batches were corrected, using the parametric empirical Bayesian framework with L/S scaling, implemented in the HarmonizR framework Version 0.0.0.9). As described by Voss & Schlumbohm et al. (2022)^27^, mean subtraction across rows was applied to batch-effect corrected TMT reporter intensities to mimic SILAC ratios, prior to data integration. Log2 transformed super SILAC ratios were median normalized across columns prior to data integration.

Processed data from individual studies was combined based on the UniProt identifier. The resulting combined dataset was subjected to HarmonizR (Version 0.0.0.9). Batch effect correction between individual datasets was performed trough the L/S scaling-based parametric ComBat mode^86^. Combined, harmonized protein abundances were mean subtracted across rows. Out of 176 analyzed cases, 9 patients were excluded from further analysis, as high blood protein yields, suppressing tumor-specific signals, were detected from LC-MS/MS measurements (Supplementary table 1a).

For the validation cohort protein abundances were log2 transformed and median normalized across columns. Samples were assigned to proteome subtypes individually. Protein abundances were reduced to the 3998 proteins, considered in the main cohort. Harmonized protein abundances from the main cohort were integrated with each individual sample. Mean row normalization was performed to adjust values from validation samples to the main cohort. Pearson correlation-based hierarchical clustering, with average linkage was applied using thePerseus software (Max Plank Institute for Biochemistry, Version 1.5.8.5)^87^.

#### Normalization of N-Glycan data

N-Glycan intensities were log2 transformed and median normalized across columns to compensate for injection amount variations.

### Quantification and statistical analysis

#### Dimensionality reduction and hierarchical clustering

Nonlinear Iterative vertical Least Squares (NIPALS) PCA and hierarchical clustering were performed in the R software environment (version 4.1.3). For Principal component calculation and visualization, the mixOmics package (Version 6.19.4.)^34^ was used in Bioconductor (version 3.14). Hierarchical clustering was performed based on pheatmap package (version 1.0.12) Pearson correlation was applied as a distance metric. Ward.D linkage was used. Pairwise complete correlation was used, to enable the consideration of missing values.

#### Consensus Clustering

To determine the ideal number of clusters from combined proteome and DNA-methylation data, Consensus Clustering was applied on normalized and integrated datasets, using the ConsensusClusterPlus package (Version 1.6) ^88^, in the R software environment (version 4.1.3). In correspondence with the current maximum number of suspected MB subtypes, the number of clusters was varied from 2 to 12 and calculated with 1,000 subsamples for all combinations of two clustering methods (Hierarchical clustering (HC) and partition around medoids (PAM)) and three distance metrics (Euclidean, Spearman, Pearson). The Ward’s method was applied for linkage. Missing value tolerant pairwise complete correlation was used, to enable the consideration of missing values. For each sample, the cluster certainty was calculated by how many times under the application of different distance metrics (Euclidan, Spearman, Pearson) and clustering approaches (k-medoids, hierarchical clustering) a sample was associated with a certain cluster, while allowing a total number of six clusters.

#### Differential analysis and visualization

Statistical testing was carried out, using the Perseus software (Max Plank Institute for Biochemistry, Version 1.5.8.5)^89^. ANOVA testing was performed for the comparison across multiple subgroups/subtypes. Factors, identified with p-value <0.05 were considered statistically significant differential abundant across groups. For the identification of subtype-specific biomarkers, Students t-testing was applied (p-value <0.05, Foldchange difference > 1.5). Visualization of t-test results and abundance distributions across groups was performed in PRISM (GraphPad, Version 5) and Microsoft excel (Version 16.5.)

For proteome data, only proteins, found in at least 30% of all proteome subtypes were considered to guarantee a high statistical validity and reasonable cohort size in differential analysis.

#### Functional annotation of data sets

REACTOME-based^90^ Gene Set Enrichment Analysis was performed by using the GSEA software (version 4.1, Broad Institute, San Diego, CA, USA)^91^, 1000 permutations were used. Permutation was performed based on gene sets. A weighted enrichment statistic was applied, using the signal-to-noise ratio as a metric for ranking genes. No additional normalization was applied within GSEA. As in default mode, gene sets smaller than 15 and bigger than 500 genes were excluded from analysis. For visualization of GSEA results, the EnrichmentMap (version 3.3) ^92^ application within the Cytoscape environment (version 3.8.2) ^93^ was used. Gene sets were considered if they were identified at an FDR < 0.25 and a *p*-value < 0.1. For gene-set-similarity filtering, data set edges were set automatically. A combined Jaccard and Overlap metric was used, applying a cutoff of 0.375. For gene set clustering, AutoAnnotate (version 1.3) ^94^ was used, using the Markov cluster algorithm (MCL). The gene-set-similarity coefficient was utilized for edge weighting.

#### Survival curves

Kaplan-Meier curves were generated for the overall survival of 121 patients. All Kaplan-Meier curves and log rank test p values were generated with PRISM (GraphPad, Version 5). A conservative log-rank test (Mantel-Cox) was used for the comparison of survival curves. A significant difference between curves was assumed at a p-value < 0.05.

#### Copy number frequency plots of Proteome and DNA Methylation data

Copy number analysis was performed on samples having both methylation and proteomic data (N=115). Samples from 450K and EPIC array were read in separately. Data were read using read.metharray.sheet () and read.metharray.exp() using the minfiData package (Version 0.36.0)^78^. For normalization, preprocessIllumina normalization using MsetEx data containing control samples for normalization of 450K array data, while for EPIC array data minfidataEPIC (Version1.16.0)^78^ was used. IlluminaHumanMethylation450kanno.ilmn12.hg19 and IlluminaHumanMethylationEPICanno.ilm10b4.hg19 were used to generate the annotation files of 450K and EPIC array data respectively. Individual sample CNV plots were generated using CNV functions as mentioned in the Conumee package (Version 1.24.0). vignette, and the segmentation information from each sample was saved which was used later for generation of cumulative CNV plot using CNAppWeb tool (PMID : 31939734)^95^. The segmentation information for all samples belonging to were combined into a single file in subgroup specific manner and then read into CNAppWeb tool (cut-off >= |0.2|) for gain or loss) [CNAppWeb user guide].

Further to map the protein abundancies to each of the chromosomes, protein names were converted to their respective gene names and a column containing mapping information for these genes was added. Copynumber (Version 1.30.0) package in R was used to generate segmentation information for these proteins in an indirect manner. Segmentation data generated was read into the CNAppWeb tool and using the cut-off mentioned above was used to map the protein abundancies to respective chromosomes.

Finally, combining the segmentation information from proteome data and methylome data in subgroup specific manner, pearson correlation-based distance plot was generated.

#### Integration of Proteome and DNA Methylation data

DIABLO or multiblock sparse partial least square discriminant analysis (sPLS-DA) using latent variables method from mixOmics (Version 6.19.4)^34^ was used for integration of proteome and methylome data to understand the correlation between the two data types. The proteome data (3990 proteins,115 samples) and methylome data (10,000 differentially methylated CpG sites, 115 proteins) were pre-processed as mentioned above. Steps followed were same as explained in the mixOmics vignette. Briefly, datasets were integrated, an output variable containing information about which subgroup the samples belong to was also supplied. Each data set is broken down into components (5 components for this study) or latent variables which are associated with the data. Components were selected using 5-fold cross validation repeated 50 times and since the groups were imbalanced lowest overall error rate and centroid distance was used. For each dataset and for each component sparse DIABLO was applied which will select variables contributing maximally to the selected component. Finally, sPLS-DA was applied to the selected variables to generate the correlation circus plot (cut-off 0.7) which gives the variables that are either positively or negatively correlation with each other.

#### Global correlation of Proteome and DNA Methylation data

To check for overall correlation between the two datasets, subgroup specific (pWNT = 13, pSHHt = 29, pSHHs = 6, pG4 = 36, pG3 = 11, pG3Myc = 20) pearson correlation (cut-off 0.7) was performed between the proteome (3990 proteins and 115 samples) and methylome (381,717 probes and 115 samples) in R (Version 4.0.5. The data was subset for correlation value and matches to their respective probes using python script in anaconda JupyterLab (Version 3.0.14). All plots were generated using ggplot2 (Version 3.3.6). Further, we also checked for the global non-subgroup specific pearson correlation between the proteome and methylome data as mentioned above and using the same cut-off only this time focusing on potential biomarkers for each subgroup and their correlation with methylation probes. Scatterplots of biomarker’s protein abundance and the M-values of CpG sites of its own gene (crossing the pearson correlation cut-off of 0.7) were plotted to confirm the correlations.

#### Quantification of Immunohistochemical stainings

Immunostained tissue sections were digitalized using a Hamamatsu NanoZoomer 2.0-HT C9600 whole slide scanner (Hamamatsu Photonics, Tokyo, Japan). Slide images were exported using NDP view v2.7.43 software. Digital image analysis was performed using ImageJ/Fiji software^96^ after white balance correction in Adobe Photoshop 2022 (Adobe Inc., San Jose, USA). Tumor areas were labelled via manually drawn regions of interest (ROIs). Tissue areas not eligible for quantification (e.g. non-tumorous tissue, technical or digital artifacts) were excluded from the analysis. Total tumor tissue areas were measured in grayscale converted images via consistent global thresholding (0, 241) and subsequent pixel quantification within the ROIs. DAB-positive pixels (i.e. brown immunostaining) were quantified on a three-tiered intensity scale after application of the color deconvolution plugin. In detail, pixels were successively quantified within three distinct thresholds [0, 134 (strong/ 3+); 135, 182 (medium/2+); and 183, 203 (weak/ 1+)]. Based on the conventional Histo-score, pixel quantities of strong, medium and weak intensity were multiplied by three, two and one, respectively, and then summed up. The hereby generated score is referred to as a digital Histoscore (DH-score).

